# Tensile Force Induced Cytoskeletal Reorganization: Mechanics Before Chemistry

**DOI:** 10.1101/2020.02.03.932863

**Authors:** Xiaona Li, Qin Ni, Xiuxiu He, Jun Kong, Soon-Mi Lim, Garegin A. Papoian, Jerome P. Trzeciakowski, Andreea Trache, Yi Jiang

**Author notes:** Co-first author (Xiaona Li, Qin Ni). Department of Chemistry, Texas A&M University, College Station, TX, USA.

## Abstract

Understanding cellular remodeling in response to mechanical stimuli is a critical step in elucidating mechano-activation of biochemical signaling pathways. Experimental evidence indicates that external stress-induced subcellular adaptation is accomplished through dynamic cytoskeletal reorganization. To study the interactions between subcellular structures involved in transducing mechanical signals, we combined experimental and computational simulations to evaluate real-time mechanical adaptation of the actin cytoskeletal network. Actin cytoskeleton was imaged at the same time as an external tensile force was applied to live vascular smooth muscle cells using a fibronectin-functionalized atomic force microscope probe. In addition, we performed computational simulations of active cytoskeletal networks under a tensile external force. The experimental data and simulation results suggest that mechanical structural adaptation occurs before chemical adaptation during filament bundle formation: actin filaments first align in the direction of the external force, initializing anisotropic filament orientations, then the chemical evolution of the network follows the anisotropic structures to further develop the bundle-like geometry. This finding presents an alternative, novel explanation for the stress fiber formation and provides new insight into the mechanism of mechanotransduction.

**Author Summary:** Remodeling the cytoskeletal network in response to external force is key to mechanosensing and locomotion. Despite much focus on cytoskeletal remodeling in recent years, a comprehensive understanding of actin remodeling in real-time in cells under mechanical stimuli is still lacking. We integrated stress-induced 3D actin imaging and 3D computational simulations of actin cytoskeleton to study how the actin cytoskeleton form bundles and how these bundles evolve over time upon external tensile stress. We found a rapid actin alignment and a slower bundle evolution leading to denser bundles. Based on these results, we propose a “mechanics before chemistry” model of actin cytoskeleton remodeling under external force.

## Introduction

Cells adapt to local mechanical stresses by converting mechanical stimuli into biochemical activities that alter the cellular structure-function relationship and lead to specific responses (1–3). Cellular response to mechanical stimulation is a balance between contractile elements of the cytoskeleton, matrix stiffness, and cell-matrix adhesions(4). Although cellular mechanotransduction has been an active field of research for a number of years, the process by which transduction of external mechanical signals across the cellular cytoplasm induces cytoskeletal remodeling is not well understood. The most important question in the field of mechanobiology is ‘*how do cells sense and integrate mechanical forces at the molecular level to produce coordinated responses necessary to make decisions that change their homeostatic state?*’

Vascular smooth muscle cells (VSMCs) provide an excellent model system to study the mechanotransduction process. The mechanism by which VSMCs sense and adapt to external mechanical forces that result in cytoskeletal remodeling (6–8) is critical in understanding arterial disease pathology. *In vivo*, they sense and respond to mechanical forces generated by pulsatile blood pressure changes by altering signal transduction pathways to induce remodeling of their cytoskeleton and adhesions (5, 6). Thus, VSMCs residing in the vessel wall are subjected to axial stress and circumferential stretch (7–9). While circumferential stretch is well accepted as an important mechanical stressor (10, 11), the axial stress in the vessel wall, which can be considered as tensile force applied to cells, has been less studied(12). Moreover, VSMC cytoskeletal network response to tensile force is not well-understood (13, 14).

To address how application of external tensile stress induces adaptive cellular remodeling, we combined imaging techniques with simultaneous mechanical stimulation of single cells using fibronectin-functionalized atomic force microscope (AFM) probes(15). In anchorage-dependent cells, external mechanical forces are imposed on a pre-existing balanced force equilibrium generated by cytoskeletal tension (16–18). Thus, forces acting on a cell will induce cytoskeleton deformation throughout the cell. Our previous experiments on VSMCs suggested that cellular adaptation to the applied tensile force is a characteristic of the integrated cell system as a whole(19). We found that mechanical stimulation increases the total intensity and alignment of bundled actin filaments. Here, we build upon these results, and ask how tensile force induces cytoskeletal remodeling and the active formation of actin bundles.

Actin cytoskeleton consists of semi-flexible actin filaments, myosin motors, and crosslinking proteins. During the adaptation process, the actin cytoskeleton remodels to better sustain the external load in two ways (20–22). On the one hand, actomyosin networks crosslinked by α-actinin and other crosslinking proteins are able to adapt external forces via fast mechanical response; the mechanical stress relaxation occurs on the timescale of seconds (23–26). On the other hand, cytoskeletal reactions, such as myosin activation, continuously convert biochemical energy into mechanical force; remodeling of the actin networks takes place on the time scale of minutes (27–29). As a result of myosin dominant mechanochemical dynamics, actin networks tend to contract (30, 31). For convenience, we called this process slower chemical response in contrast to the faster mechanical response. Observation of actin filament microstructure changes under these two mechanisms requires super-resolution and ultra-fast imaging, which is not achievable using our present experimental methods. Prior computational models have investigated actin bundles generation and remodeling due to slower biochemical reactions (32–36), but how external stimuli induce the active formation of actin bundles is still poorly understood.

To better understand the detailed spatiotemporal dynamics of cytoskeletal reorganization upon the external mechanical loading, we simulated the mechanical and chemical dynamics of the actin cytoskeleton using the MEDYAN (MEchanochemical DYnamics of Active Network) package(37). In our simulations, we model the active cytoskeletal networks using polymer mechanics of semi-flexible filaments, crosslinking proteins, and motor proteins. A stochastic reaction-diffusion scheme was used to simulate biochemical reactions, including myosin activation, crosslinking proteins binding, and actin filament assembly. Additionally, we applied tensile external force to the actin network to mimic the AFM mechanical stimulation experiments. In these simulations, a few filaments anchored to a pseudo-AFM probe were initialized in addition to a free filament pool. The external force was applied via moving the pseudo-AFM probe, and the amplitude of z-axis displacement directly determined the magnitude of the force. In highly crosslinked actomyosin networks, the external force exerted on a small fraction of filaments would transmit to the entire filament network that changes its homeostatic state in microseconds(38): this will be considered as the fast mechanical response. After each tensile force was applied, the system was allowed to evolve for minutes, such that we can study how the actin network transforms under the slower chemical response.

In the present work, to investigate the detailed spatiotemporal cytoskeletal remodeling during tensile mechanical loading, we integrated experimental and modeling approaches. Both experiments and simulations suggest that the external tensile force applied on actin networks quickly induces alignment of actin filaments along the direction of force, and this directional alignment is independent of longer timescale biochemical responses. We also observed filament bundle formation as a result of external tensile force; however, the bundle development relies on both the faster mechanical response and the slower chemical response. We hypothesized that the cellular cytoskeletal adaptation to external mechanical forces and filament bundle formation follows a “mechanics before chemistry” process.

## Results

### Actin cytoskeleton reorganization of live VSMCs under mechanical stimulation reveals two types of responses

Live VSMCs were subjected to the mechanical loading delivered by the AFM probe at the apical cell surface (Figure 1a). Vertical forces (along the z-axis) applied through a fibronectin (FN) functionalized probe induces cytoskeletal remodeling by pulling on cortical actin through a FN-integrin-actin linkage (9, 19). Cell responses to the probe displacement over time were recorded using spinning-disk confocal microscopy. The reconstructed 3D-images of the actin cytoskeleton were used to segment actin bundles in 3D (Figures 1b & 1c). We used these 3D bundles to calculate an average fiber alignment index in the direction of the pulling force. The alignment index is defined as the average of *cos(θ)*, where *θ* is the acute angle between each filament segment and the direction of the force (Z-axis) (Equation 1 in Methods). With a value between 0 and 1, the alignment index equals to 1 for perfect alignment with the Z-axis, 0.5 for completely random, and 0 for alignment perpendicular to the Z-axis. The alignment index increases right after the application of an external force, but levels off (Figure 1d) upon larger pulling forces. Note that the small alignment index value is because of the cells initially lying flat on the substrate, and the majority of the filaments were perpendicular to the Z-axis. In addition, the normalized fluorescence intensity of actin-mRFP, which corresponds to the total amount of measurable actin bundles, increased steadily as the AFM displacement continued (Figure 1d). These experimental results show a force-induced actin cytoskeleton remodeling via the directional alignment and stress fiber consolidation.

**Figure 1.**
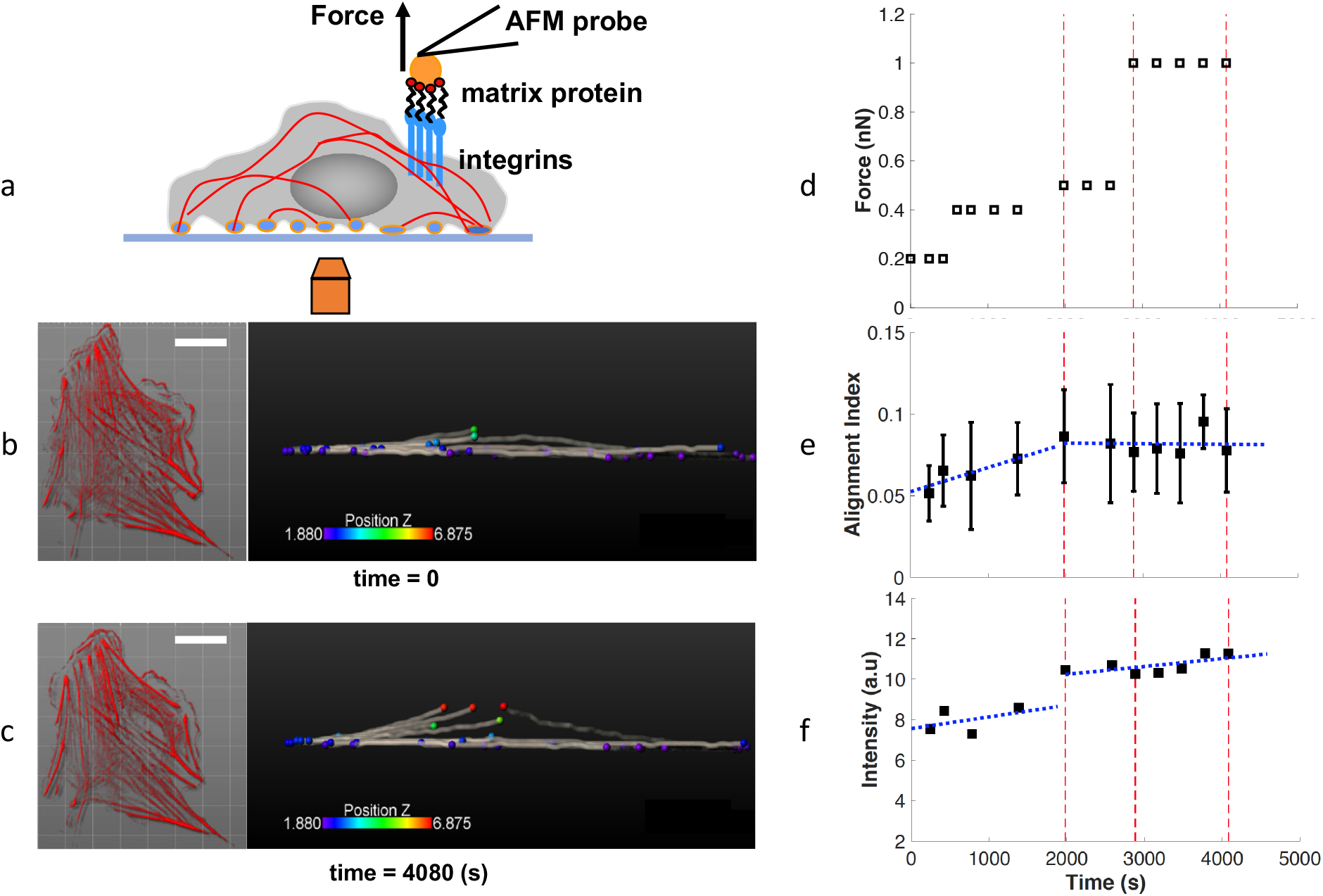
Response of VSMC to external pulling forces, (a) Schematic of a VSMC were mechanically stimulated with a fibronectin functionalized AFM probe and simultaneously imaged by spinning-disk confocal microscopy. (b) Fluorescence images of VSMC expressing actin-mRFP (left) and the 3D reconstructed image of the representative segmentation of actin filament bundles (right) for before the AFM probe displacement at time 0 min, and after the AFM probe displacement at time 68 min (c). Scale bar: 20 μm. (d) The scheduled pulling force in three phases: small, intermediate and large forces. (e) The alignment index for the actin filament bundles increased rapidly as small force was applied, but slowed down as the force increased. (f) The normalized intensity for actin-mRFP increased steadily through all force ranges. Blue lines: piece-wise linear fit.

### Rapid formation of actin bundles in response to tensile force in MEDYAN simulations

To understand the experimental observations of actin alignment and intensity change (Figures 1e & 1f), and explore the molecular details of the actin cytoskeleton reorganization, we designed MEDYAN simulations of actin networks with external tensile force, mimicking the AFM experiments on VSMCs. We generated 300 free filaments in a 3×3×1.25 μm^3^ simulation box, initially as a random network, and additional 30 filaments attached to an AFM probe located at the center of the upper boundary of the simulation box. The simulation box contains 20 μM of actin, 0.1 μM of myosin, and 2 μM of a-actinin crosslinkers. The simulated AFM probe was displaced by a distance *d*, every 150 seconds. Each pulling created a 250 nm or 500 nm step displacement on the AFM tips, generating tensile force to AFM-attached filaments via stiff harmonic springs (Figure 2a). The amplitude of step displacement size *d* is linearly proportional to the pulling force of the AFM probe. Chemical interactions, including filaments treadmilling, myosin activation, and α-actinin crosslinking, take place throughout the simulations. We varied the pulling patterns (Figure 2b) to simulate the different pulling forces in the experiment (Figure 1d).

**Figure 2.**
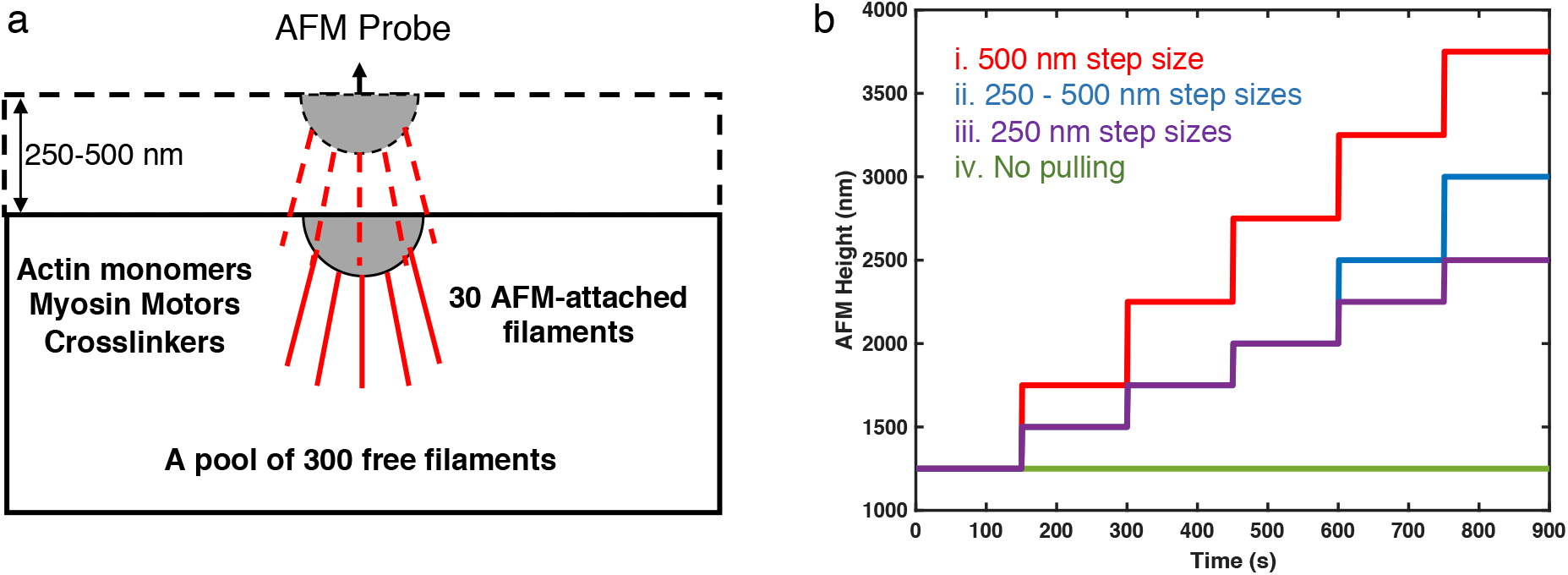
(a) A sketch of the simulation setup. The simulation box is 3 μm in x and y directions, and the initial height (z-direction) is 1.25 μm. The simulation box contains 300 free actin filaments, as well as diffusible G-actin, myosin, and α-actinin linkers. A semi-spherical AFM tip is located on the top of the upper boundary, and 30 filaments are attached to the tip via stiff harmonic springs. (b) AFM probe position, equivalent to the height of upper boundary, as a function of time for Cases i-iii. The control case (Case iv) is without AFM-probe and no filament attachment, with only the upper boundary moving in the same way as in Case i to avoid potential boundary effects.

Interestingly, pulling on only a small fraction of AFM-attached filaments is sufficient to alter the actin filament structure of the entire network. After 900s and five AFM probe pulling steps, each with *d*=500 nm, the actin networks reorganized into a bundle (Figure 3a, and Video 1), which is approximately 2 μm long and around 500 nm thick under Case i pulling pattern (red line in Figure 2b). These actin bundles have mixed filament polarity, i.e., plus ends or minus ends of filaments are randomly distributed (Figure S1 in Supporting Information), implying that they are closer to stress fibers rather than unipolar actin filaments in filopodia (21, 39). In contrast, actin networks free of external force geometrically collapsed into a globular cluster-like structure (Figure 3b, and Video 2), as a result of myosin-driven contractility.

**Figure 3.**
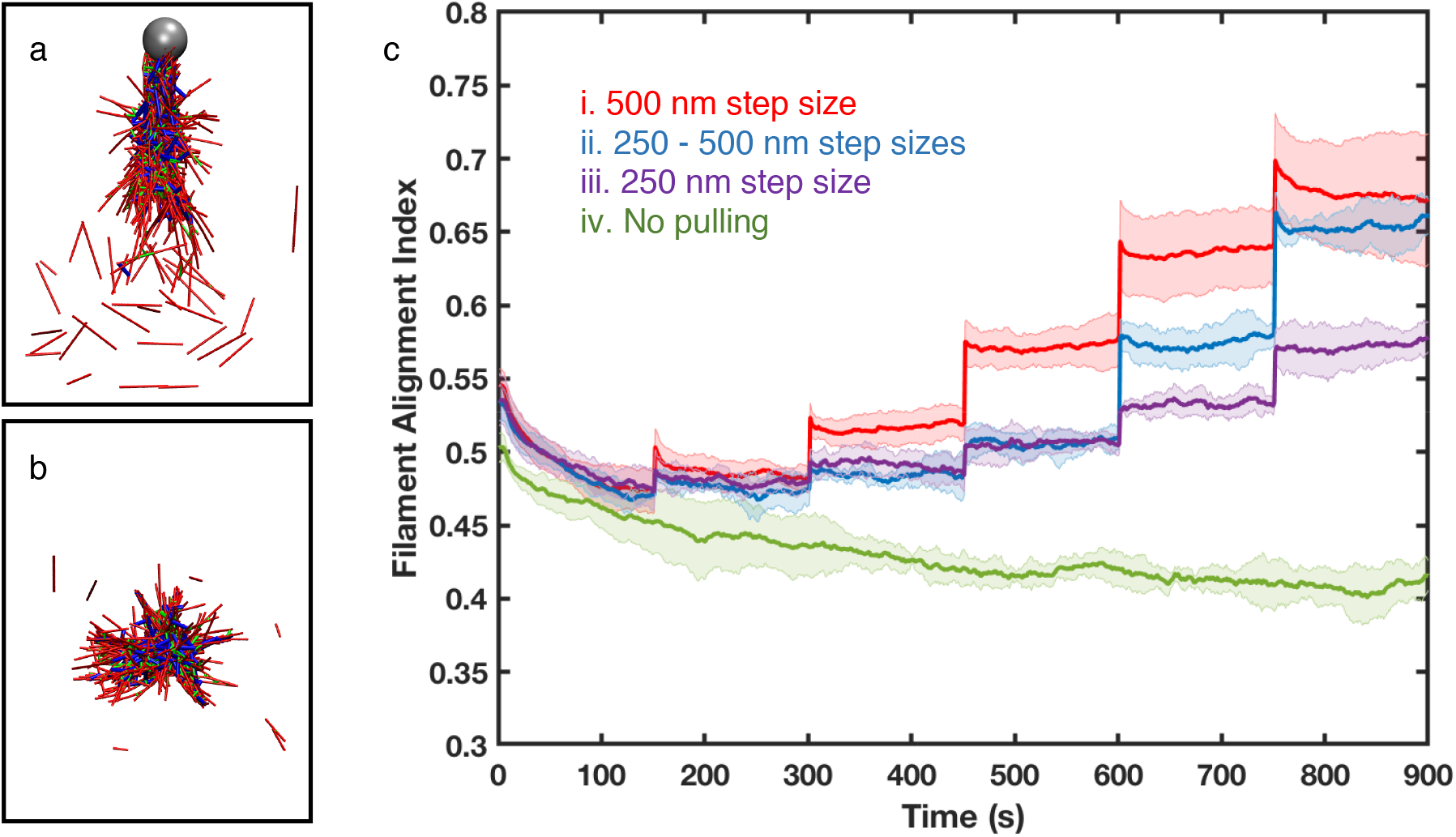
(a-b) Representative snapshots of (a) bundle-like actin networks under Case i pulling condition at time = 900 s, and (b) cluster-like actin networks without external force at time = 900 s. Actin filaments, myosin motors, and crosslinkers are shown in red, blue, and green cylinders, respectively. The grey sphere in (a) represents the AFM probe. (c) Filament alignment index along the Z-axis for 500 nm step size (red, Case i), mixed step sizes (250 nm for the first three pulling events and 500 nm for the last two, blue), 250 nm step size (purple, Case ii) and no AFM pulling (green, Case iii). α-actinin linker:actin is 0.1 and myosin:actin is 0.005 in all simulations. Error bars represent the standard deviation from the mean in 5-10 replicate simulations.

### Fast tensile force-induced actin alignment

To investigate the development of actin filament alignment during actin bundle formation, we calculated the alignment index *cos(θ)* as described in the experimental section. The alignment index increases immediately after each of the AFM pulling events in all three pulling patterns tested (Figure 4c, Case i-iii). In the simulation, the mechanical equilibration is instant, therefore these rapid jumps suggest very strong mechanical responses. Moreover, the directional filament alignment is directly regulated by the strength of the external tensile force, since reducing the pulling step size amplitude (compared to Cases ii and iii) results in a weaker alignment response. On the other hand, the directional alignment barely changes at long timescale in all step size patterns. Since the long timescale response is regulated by slower chemical evolutions, we hypothesize that filaments directionally aligned in response to tensile force is primarily due to fast mechanical adaptation.

**Figure 4.**
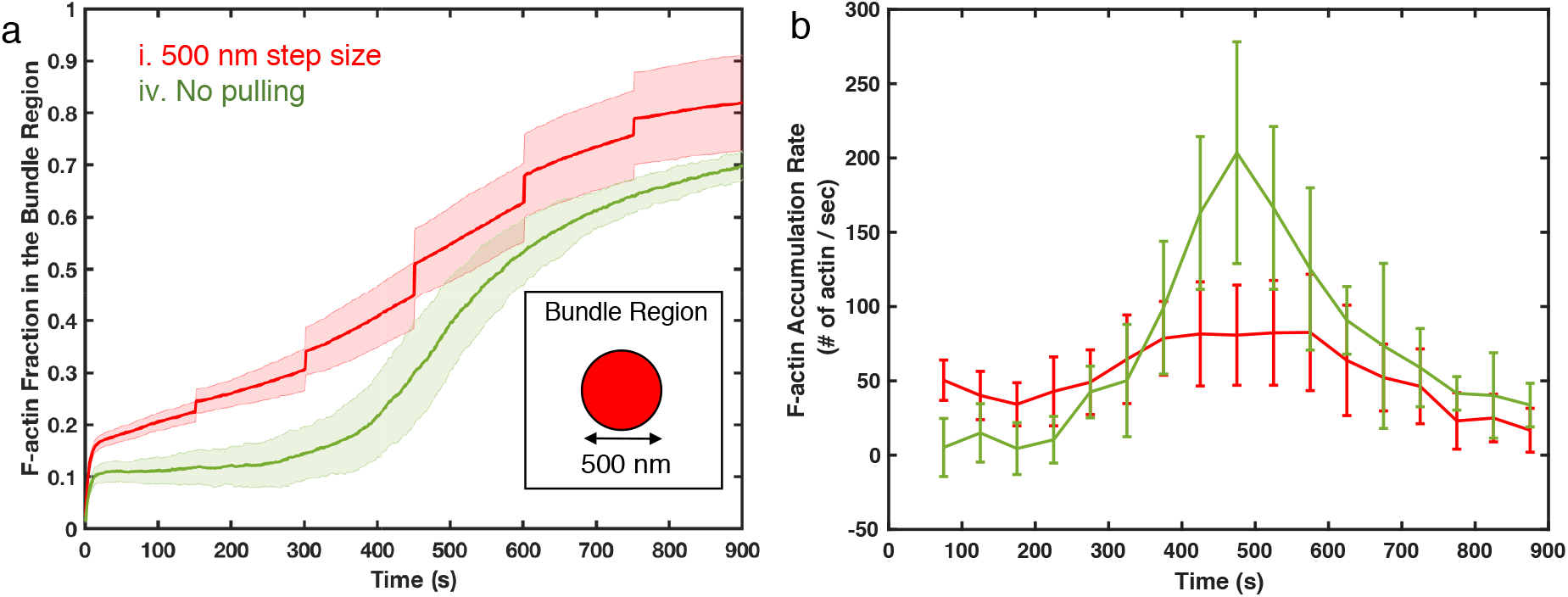
(a) The F-actin fraction in the bundle region as a function of time of networks under 500 nm displacement (red, Case i) and networks free of external force (green, Case iv). The bundle region is defined as the volume under the AFM probe, which is a cylindrical region of 500 nm in diameter and the height of the simulation box, which varies over time based on the position of the AFM probe. Insert shows a 2D illustration of the bundle region. (b) The rate of F-actin accumulation in the bundle region from simulations with the AFM probe pulling (red, Case i) and without AFM pulling (green, control Case iv). The accumulation rates are calculated by linear-fitting of the data points every 50 seconds. (a-b) Shaded colors and error bars are the standard deviations of 10 replica simulations for Case i and 5 replicas for Case iv, respectively.

### Two-step development of actin bundles relies on both faster mechanical alignment and slower chemical stabilization

To further analyze formation and evolution of actin bundles, we next defined a cylinder-shaped boundary under the AFM probe (500 nm in diameter). More than 80% of the total F-actin accumulated within this boundary towards the end of simulations under the Case i pulling condition, suggesting that monitoring the F-actin’s accumulation in the bundle region provides a simple but robust way to quantify the bundle development process. We observed instant F-actin accumulation after each AFM pulling event (Figure 4a), while reducing step size hindered the accumulation (Figure S2 in Supporting Information). Similar to the directional alignment, these results suggest that actin bundle development relies on the fast mechanical response.

Somewhat surprisingly, the accumulation of actin filament into the bundle keeps increasing steadily between pulling events, suggesting that slower chemical dynamics contribute to bundle development. To capture the long timescale of F-actin accumulation, we calculate the F-actin accumulation rate in the defined bundle region (Figure 4b). The control case without external pulling (iv, green line) shows the biochemically driven F-actin accumulation, as a result of myosin-induced contractility. Similarly, the accumulation rate of F-actin during the intervals between pulling (Case i, red line) is always positive, showing a net accumulation of F-actin. The rate of F-actin accumulation for bundling is lower than that for contractile clustering.

To further explore the significance of chemical evolution in bundle development, we tested three different conditions with “insufficient” chemical evolution. First, we reduced the myosin concentration from 0.1 μM to 0.02 μM. Without sufficient myosin, the network was unable to generate enough contractility to integrate the network, leading to high actin filament dispersion (Video 3). Second, by reducing a-actinin crosslinker concentration from 2 μM to 0.4 μM, the F-actin network could not form properly (Video 4). Although myosin motors still generated contractility, the fiber network is fragmented as disconnected actin foci. Lastly, we shorten the time between each pulling from 150 seconds to 10 seconds. Only a small fraction of filaments bundled together and followed the upward movement of the AFM probe, disconnecting from the rest of the filaments (Video 5).

F-actin distribution further showed that the tensile AFM pulling immediately stretches the actin fiber network along the direction of pulling force (Figures 5a-c), leading to a wider distribution. As a result, the standard deviations (σ) of these distributions increased right after pulling (Figure 5e). When we measure the radius of gyration (Rg) to quantify the cluster size of actin networks, we also find instantaneous jumps similar to those in the filament alignment and accumulation results (Figure 5f). These instant stretches eventually shape actin networks into the thinner bundles.

**Figure 5.**
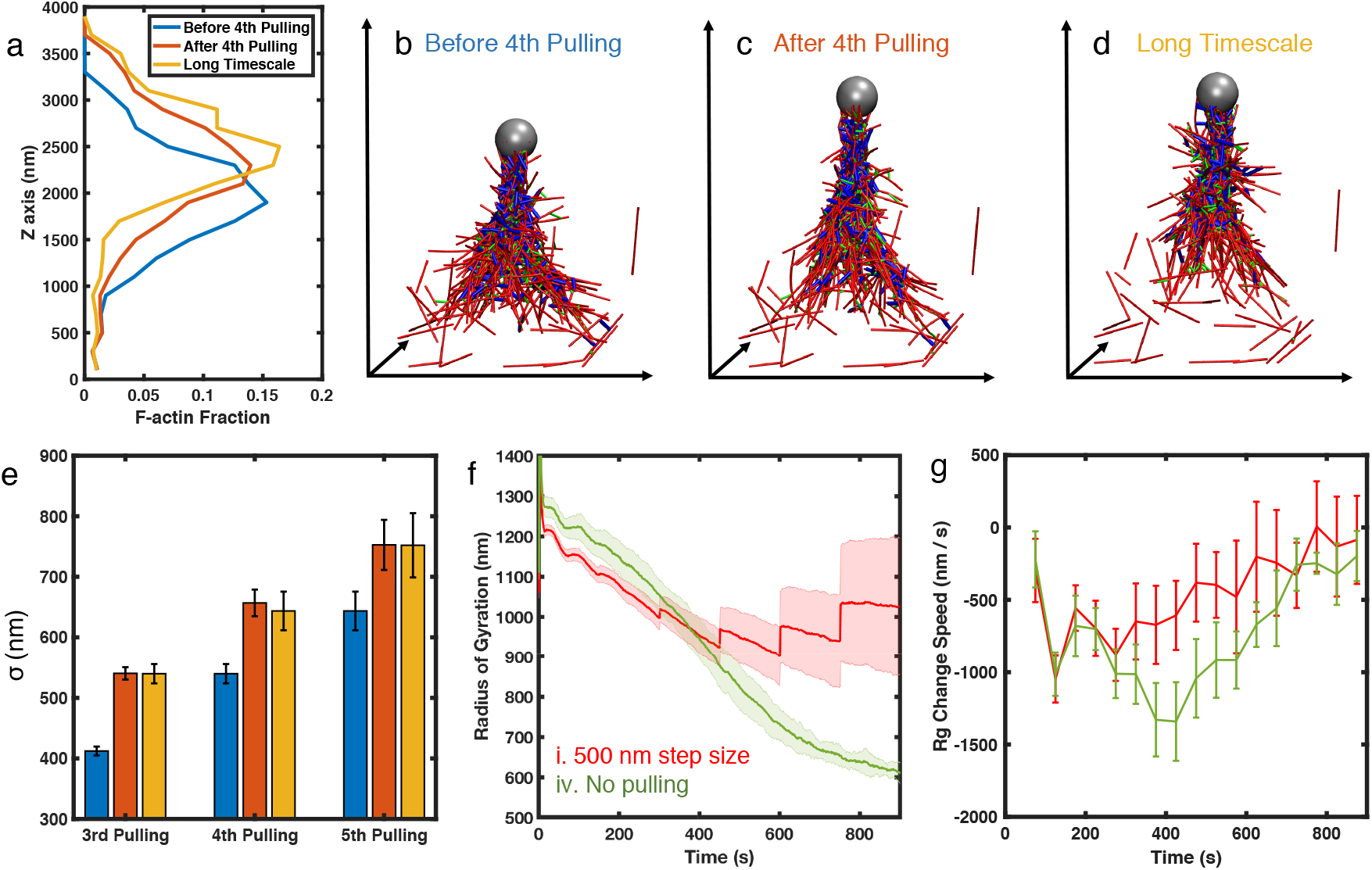
(a) F-actin distribution along the force direction (Z-axis) of the representative trajectory before the 4^th^ pulling at time = 600 s (blue), after the 4^th^ pulling at time = 601 s (orange), and after the long-timescale chemical evolution at time = 750 s (yellow). (b-d) Corresponding simulation snapshots before pulling, after pulling, and after chemical evolution, respectively. (e) Standard deviations (σ) of the F-actin distribution along the force direction before pulling (blue), after pulling (orange), and after 150 seconds of chemical evolution (yellow) at the third, fourth, and fifth pulling events. σ are averaged over 10 duplicated trajectories, and error bars represent the standard errors. (f-g) The radius of gyration (f) and the change speed of Rg (g) of networks under 500 nm AFM step size (red) and without AFM pulling (green). Shaded colors and error bars are the standard deviations of 10 duplicated trajectories for Case (i) and 5 duplicated trajectories for Case iv.

Furthermore, these actin bundles maintain their geometric structures at a longer timescale. The F-actin distribution of bundle networks shifts slightly towards the force direction after 150 seconds of chemical evolution (Figure 5a and 5d), but the shape and the standard deviation from the mean, σ, remain almost the same (Figure 5e). In addition, the contraction rate, measured as Rg change speed, is much slower than that for the cluster networks under no AFM pulling (Figures 5g). These observations are consistent with the slower F-actin accumulation rate in the bundle region, as shown in Figure 4b, suggesting that the bundle structure is more stable than the cluster induced by myosin. These results are also in agreement with the fact that the actin bundle can preserve its shape and would not contract into clusters under myosin driven contractility at long timescale.

## DISCUSSIONS

Mechanotransduction is the process by which cells convert mechanical stimuli into biochemical activity. A key aspect of the mechanotransduction is that cells remodel their cytoskeleton in response to mechanical stimuli. To study external force-induced adaption of the actin cytoskeleton, AFM was used to apply external forces on single cells adherent on a substrate. Cell responses measured through probe displacement over time are directly dependent on intrinsic contractility that modulates the function of the actomyosin apparatus. The observed rapid rise in actin fiber alignment upon tensile force stimulation contrasts with the steady increase of actin fluorescence intensity, leading to our hypothesis of ‘mechanics before chemistry’: fast mechanical stimulation-induced actin bundle alignment, followed by a slower chemical driven process to stabilize the actin bundle structure.

To explore this hypothesis, we developed a new capability in the MEDYAN software that mimics the conditions of our AFM experiments. Our simulation results reveal that tensile force triggers a rapid mechanical adaptation of actin networks that forces filaments to align along the pulling direction and encourages actin bundle (stress fiber) formation. We also found that slower biochemical evolution is essential to the formation of stress fibers, which requires integration of actin networks through α-actinin crosslinking followed by myosin activation and eventual further actin recruitment to the stress fiber. Moreover, we found that actin bundles generated in our simulations are stable since they contract much slower than networks free of external force.

Thus, our simulations agree with the experiments, supporting a “mechanics before chemistry” hypothesis as an alternative, novel explanation regarding how active cytoskeletal networks adapt to external mechanical stimuli in real-time. In the control case of actin networks without external forces, primarily driven by myosin motors, actin network contraction does not have a bias towards a specific direction, leading to an isotopically collapse into globular clusters (Figure 6a). The external tensile force first stretches the actin cytoskeletal network, forcing filaments to align, as a rapid mechanical response, which initializes anisotropic bundle-like structures. Longer time scale chemical processes further stabilize the bundle structures that can preserve the anisotropy (Figure 6b). As a result, the contractility generated by subsequent chemical evolution follows the anisotropic distribution, which strengthens actin bundles by recruiting more actin filaments while maintaining the bundle shape.

**Figure 6.**
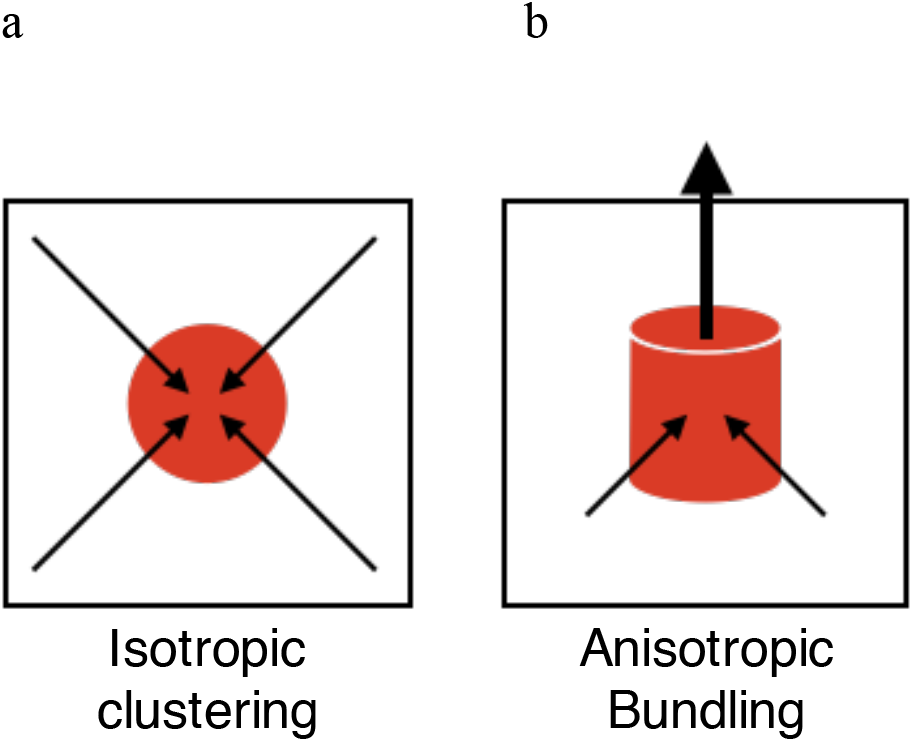
Motor-driven biochemical evolutions generate contractility that induces the geometric collapse of the actin network. In random networks without external forces, the geometric collapse would be isotropic, causing filaments to cluster into globular foci (left). However, the external tensile force inducing filament directional alignment that favors anisotropic biochemical evolutions, results in filament bundling.

Stress fibers play a crucial role in maintaining cellular shape and supporting force transmission to and from extracellular substrates. Numerous studies have demonstrated the direct coupling between mechanical forces and biochemical signaling. For example, stretching of specific focal adhesion proteins can expose either cryptic binding sites or cryptic phosphorylation sites, thus triggering signaling pathways (40, 41). Moreover, responses to mechanical forces induced by changes in substrate stiffness will further trigger the adaptation of their cytoskeletal network in less than 100 ms(42), proposing a ‘mechanics first’ mechanism of cellular response that supports our hypothesis. Thus, when the cell experiences an external force, the cytoskeletal adaptation will first elicit the actin fiber rearrangements (mechanical) before spending ATP to initiate the biochemical reactions (chemical).

In summary, we integrated *in vitro* and *in silico* modeling to investigate the effects of external load on the cytoskeleton network. Both experimental and simulation results suggest that tensile stress aligns the fibers along the direction of the external stress, before biochemical evolution to further remodel the network. This result suggests the short timescale mechanical structural adaptation operates before slower biochemical processes, which can have important implications to mechano-signal transduction.

## Methods

### Experimental Methods

#### Vascular smooth muscle cell culture and transient transfections

VSMC were previously isolated from rat cremaster arterioles(43) and handled as previously described(44). Briefly, a smooth muscle cell culture media containing Dulbecco’s Modified Eagle Medium (DMEM) supplemented with 10% fetal bovine serum (FBS), 10 mM HEPES (Sigma, St. Louis, MO), 2 mM L-glutamine, 1 mM sodium pyruvate, 100 U/ml penicillin, 100 μg/ml streptomycin and 0.25 μg/ml amphotericin B. Cell cultures were trypsinized and transient transfections for actin-mRFP were performed according to manufacturer’s protocol by using the Nucleofector apparatus (Lonza, formerly Amaxa Biosystems, Gaithersburg, MD) with Nucleofector kit VPI-1004. Transfected cells were plated on 60 mm MatTek glass bottom dishes (Ashland, MA, USA) in phenol-red free cell culture media, and incubated overnight in 5% CO2 at 37 ^°^C. Actin-mRFP plasmid was a gift of Michael Davidson (Florida State University, Tallahassee, FL). Unless otherwise specified, all reagents were purchased from Invitrogen (Carlsbad, CA, USA).

#### Vascular smooth muscle cell imaging

The integrated microscope system used for these studies was described in detail(45). Briefly, an atomic force microscope (XZ Hybrid Head, Bruker Instruments, Santa Barbara, CA) was combined with a Yokogawa CSU 22 spinning-disk confocal microscope to enable mechanical stimulation of live cells and simultaneous visualization of molecular dynamic events at the subcellular level in real-time. A PLAN APO TIRF 60x oil 1.45 NA objective lens was used for imaging live cells expressing fluorescent protein constructs. Confocal images were acquired as 3D stacks of 20 planes at a 0.25 μm step size with an exposure time of 100 ms.

#### AFM mechanical stimulation to VSMCs

Tensile stress was applied to live VSMCs using an atomic force microscope probe functionalized with a 2-micrometer glass bead coated with fibronectin (Novascan Technologies, IA, USA)(9). Formation of a functional linkage between the AFM probe functionalized with fibronectin and cortical cytoskeleton via integrins enables mechanical stimulation of the cell through the application of tensile forces. A mechanical stimulation experiment consists in four segments of force application. First, the probe is brought in contact with the cell for 20min to allow the formation of a functional adhesion through recruitment of integrins and focal adhesion proteins. During this time, the probe rest on the cell surface, and no tensile force is applied. The second step consists in the application of small tensile forces (i.e., mechanicals stimulation by 0.2-0.4 nN) to further reinforce the adhesion by enhancing protein recruitment at the respective site. Then, the mechanical stimulation of the cell with low (~0.5 nN) and high (~1 nN) magnitude forces consisted of controlled upward movement of the cantilever in discrete steps at every 3–5 minute intervals. The same force regime mechanical stimulation was applied for 20–25 minutes each, while the actin cytoskeleton was imaged by spinning-disk confocal microscopy after each force application(9). The AFM data were acquired using NanoScope (8.15, Bruker) Software and were processed off-line in MATLAB (R2019b, Mathworks) and Excel (Microsoft).

#### Three-dimensional image analysis

For each raw three-dimensional (3D) image volume at a specific time point, imaging data in z-direction were interpolated by linear interpolation to generate a new sequence. Spatial sizes of a voxel in three dimensions were not all equal, i.e., Δ*x* = Δ*y* = 0.178 *μm*, and Δ*z* = 0.25 *μm*. The resulting image sequences were imported to Imaris (v.9.3.0, Oxford Instruments, Inc.) for Automatic Tracing analysis. The coordinates of branch points from this tracing analysis was exported and saved in result data files. The 3D coordinates of all paired points that are 10 points apart along a given trace are used to compute the alignment index:

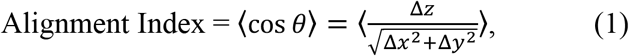

where Δ*x*, Δ*y*, Δ*z* are differences of the paired points in x, y, and z direction, respectively. The resulting set of measurements along each trace was averaged as an estimate for the angle between each trace and the z-axis. As an aggregated measure for trace angles at each time point, angle measures from all traces at a given time point were further averaged.

### Simulation Methods

A computational model for mechanochemical dynamics of active networks (MEDYAN)(37) was used to simulate the actin cytoskeletal network with an external pulling force. In this model, actin filaments are considered as connected “cylinders” with strong axial stretching stiffness. The cylinder itself is unbendable, and the radial deformation of filaments is realized by bending between two neighboring connected cylinders. Each cylinder consists of up to 40 actin monomers, where a full cylinder is 108 nm long and has 4 possible binding sites for myosin motors and crosslinkers. Myosin motors are modeled as harmonic springs that can walk towards filament plus end with equilibrium length from 175 nm to 225 nm based on the non-muscle myosin II. Crosslinking proteins are also modeled as harmonic springs with an equilibrium length for α-actinin (30-40 nm). The main chemical events we considered in this work include filament polymerization and depolymerization, binding and unbinding of myosin and crosslinker, and myosin activation. These reactions are mechanochemically sensitive and are modeled by an efficient Next Reaction Method based on the Gillespie algorithm(46, 47). Simulation parameters and other model details can be found in Supplementary Information and a previous publication(37).

We initialized a 3 ×3 × 1.25 μm^3^ simulation volume with a 250 nm semi-spherical AFM tip that attached to the upper boundary. At time 0 sec, 300 seed filaments, each with 40 monomers, were randomly created in the network, defined as the free filament pool. These filaments free from AFM attachment are allowed to polymerize and depolymerize on either the plus end or the minus end. To appropriately transmit the external force generated by AFM displacement to the actin network, another 30 seed filaments with their minus-end attached to the AFM tip via stiff harmonic springs were initialized (Figure 2a). Only plus ends of these filaments are allowing to polymerize and depolymerize. At the start of simulations, free G-actin was added to the network to ensure the total actin concentration is 20 μM. Since the concentration is much larger than the critical concentration(48), seed filaments would grow rapidly and reach an average F-actin length of ~0.8 μm in a few seconds of simulation. 0.1 μM myosin motors and 2 μM α-actinin crosslinkers were added after 5 seconds of simulation. The addition of myosin and α-actinin linkers connect the free filament pool to AFM-attached filaments while generating contractility to allow the network to self-construct.

The external tensile force from the AFM tip was implemented as follows. The network was allowed to evolve for 150 seconds before AFM probe vertical displacement (i.e., vertical pulling). Each probe displacement created a 250 nm or 500 nm step displacement of the AFM tips, generating tensile force to AFM-attached filaments via stiff harmonic springs. To ensure the energy was properly minimized, each displacement step was broken up into 100 sub-steps (2.5 nm or 5 nm displacement per 0.01 s). Networks were mechanically equilibrated after each sub-step, and displacement would create additional simulation space by raising the upper boundary. Since all AFM probe displacements were finished in 1s and each mechanical minimization was instant in the simulation, we are able to treat the network change before and after displacement as a fast mechanical response that is independent of biochemistry. Networks were allowed to evolve for another 150 seconds before the next AFM displacement step (Fig. 2b). During the 150 second period, cytoskeletal network remodeling was biochemically dominated by filament treadmilling, myosin activation, and α-actinin linker binding and unbinding. Since the time interval between two displacement steps is much longer than the pulling time (1 second), we define the network evolution during each 150 seconds as the long timescale biochemical response. We applied the tensile AFM step displacement 5 times during each simulation, taking 900 seconds of simulation time in total.

The present work tested four different tensile force conditions. For convenience, we labeled them as Case i-iv in decreasing order of displacement sizes (Fig. 2b). In Case i, a constant 500 nm step size was applied. This step size exerted an instantaneous force on the AFM attached filaments twice higher than the 250 nm step size. In Case ii, we used mixed step sizes: in the first three pulling events, each step generates 250 nm displacement, and in the last two pulling events, each step generates 500 nm displacement. In Case iii, we reduced the displacement size to constant 250 nm, implying a weaker external force. In the last case, we did not apply any external force to the network, hence, all 330 filaments were in the free filament pool. However, the upper boundary in Case iv would still move up in the same way as for Cases i-iii to avoid any influences from the boundary effects.

## Acknowledgements

This work was supported, in part, by Public Health Service grants R01CA201340 and 1R01EY028450 from the NIH/NCI and NIH/NEI, respectively (to Y.J.), K25CA181503 and U01CA242936 from NIH/NCI (to JK), National Science Foundation grant CHE-1800418 (to GP). The experimental work was supported by NSF CAREER 0747334 award to AT.

## Supplementary Figure

**Figure S1.**
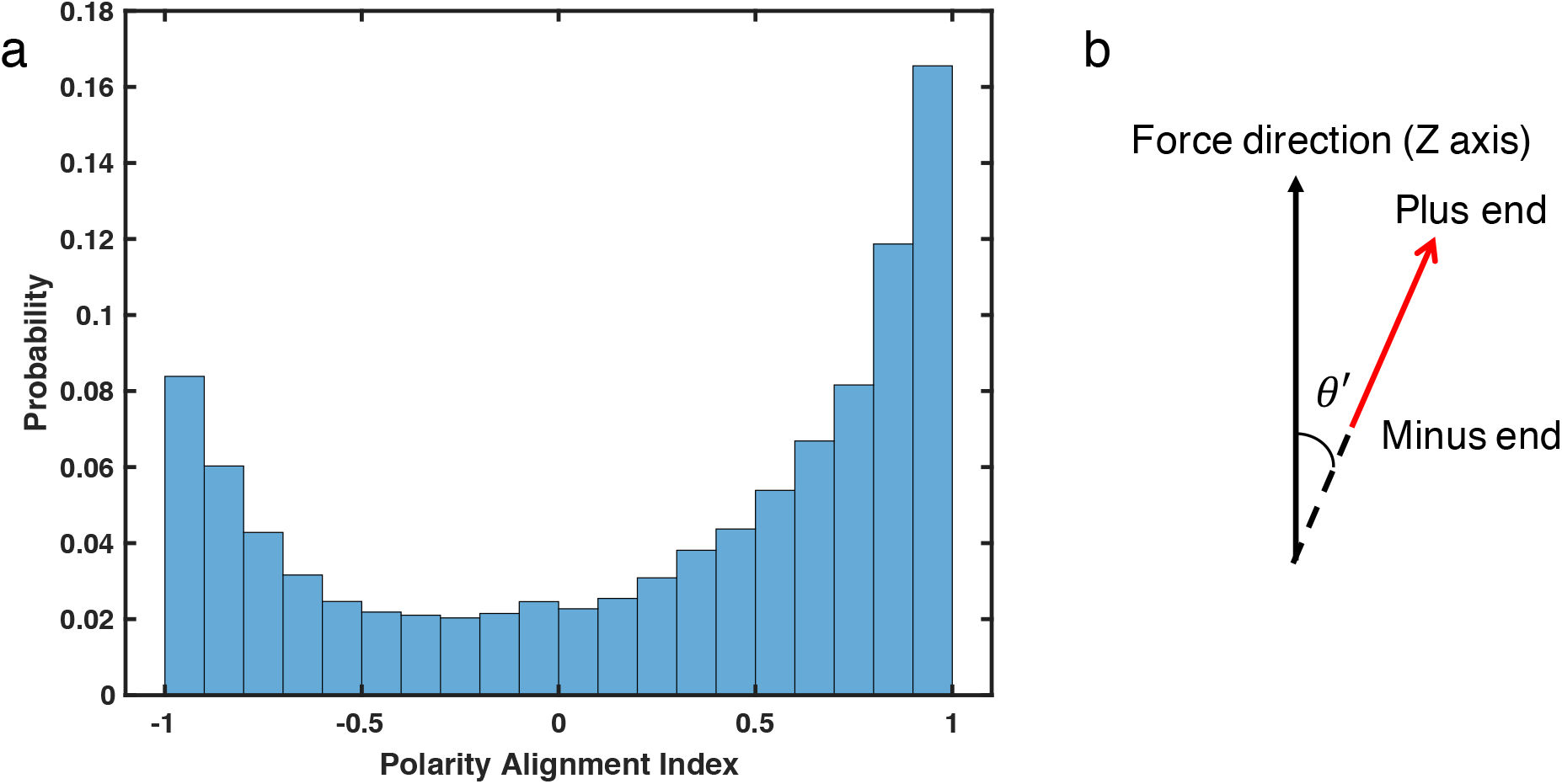
(a) The probability distribution of filament polarity alignment index for bundle-like networks under pulling condition Case i. Data are taken from 751s~900s out of 5 duplicated trajectories. (b) The polarity alignment index is defined as cos′ *θ*, where θ′ is the angle between a filament vector and the force direction. The filament vector (red arrow) in this case, considers the polarity of plus end and minus end. (a-b) The distribution spreads across [-1,1], suggesting actin bundles generated in this work is similar to stress fiber with mixed polarity. For a filopodia-like bundle, where actin fibers have the same polarity, the alignment index should have a single peak at either 1 or −1.

**Figure S2.**
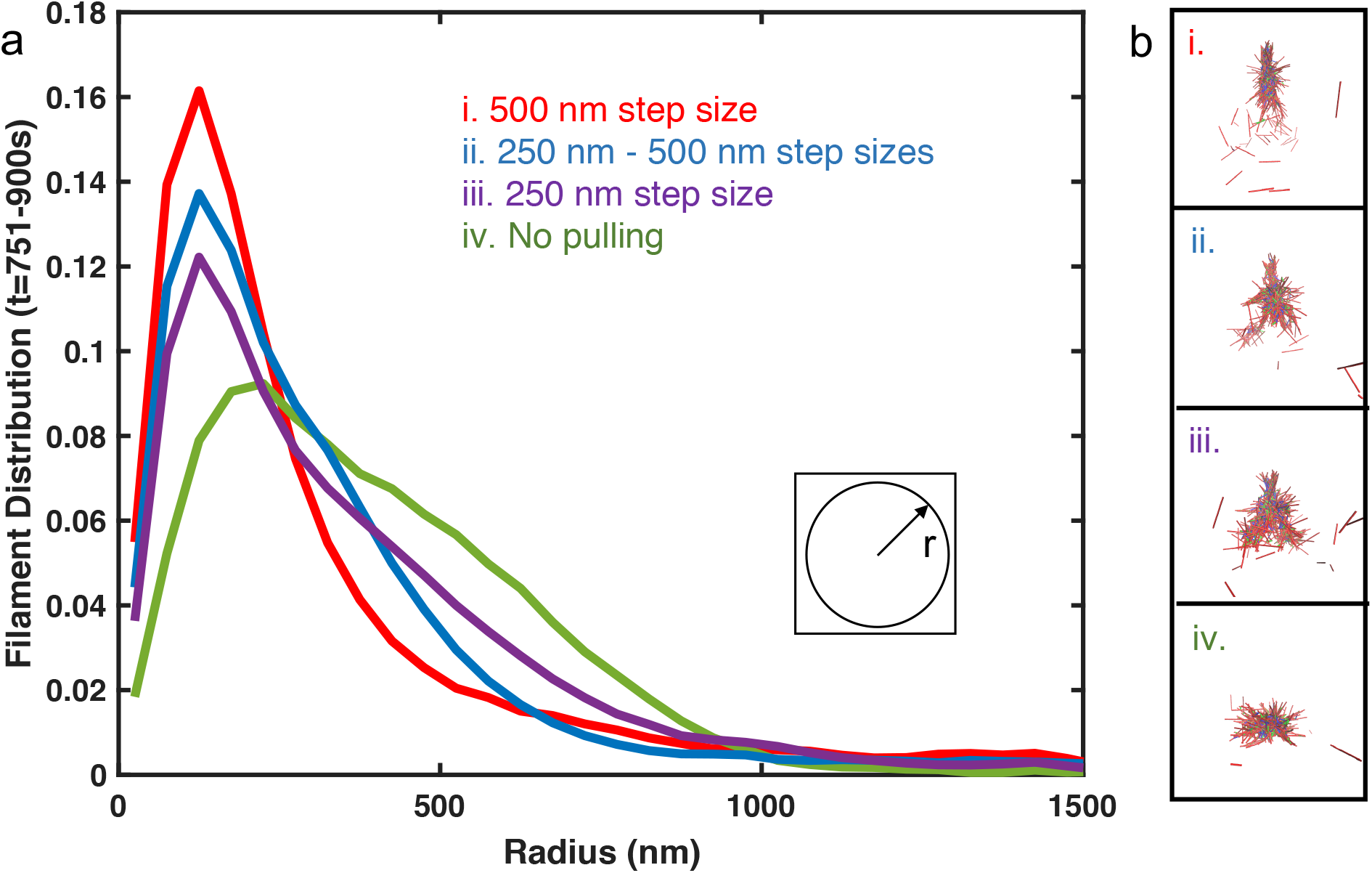
(a) F-actin radial distribution after the last pulling event (751~900s) under pulling condition Case i-iv. (b) The most representative snapshots at t = 900s for each case.

## Supplementary Videos

Video 1. Actin filament bundle formation under tensile force induce by pseudo-AFM probe with step size d=500nm (Case i pulling condition). The network contains 330 filaments with 30 filaments attached to the pseudo-AFM probe. The grey sphere represents the pseudo-AFM probe, and red, blue, and green cylinders represent the actin filaments, crosslinkers, and myosin motor mini filaments, respectively. C_actin_ = 20 μM, C_NMII_ = 0.1 μM, and C_crosslinkers_ 2 μM.

Video 2. Actin network geometrically contracts into the cluster-like structure without external force. The network also contains 330 filaments, and no filaments are attached to the pseudo-AFM probe. Red, blue, and green cylinders represent the actin filaments, crosslinkers, and myosin, respectively. C_actin_ = 20 μM, C_NMII_ = 0.1 μM, and C_crosslinkers_ = 2 μM.

Video 3. Actin network evolution under Case i pulling condition (d=500nm), but myosin concentration is reduced to 0.02 μM. Under this condition, the network does not contract, and the majority of the network remains random throughout the simulation. The grey sphere represents the pseudo-AFM probe, and red, blue, and green cylinders represent the actin filaments, crosslinkers, and myosin, respectively. C_actin_ = 20 μM, and C_crosslinkers_ = 2 μM.

Video 4. Actin network evolution under Case i pulling condition (d=500nm) with lower crosslinker concentration (C_crosslinkers_ = 0.4 μM). Although the network still contracts, the AFM attached filaments disconnected from the free actin filament pool after ~ 300s. Eventually, the networks become a small filament bundle attaching to AFM probe at the top of the network and a disconnected larger cluster at the bottom. The grey sphere represents the pseudo-AFM probe, and red, blue, and green cylinders represent the actin filaments, crosslinkers, and myosin, respectively. C_actin_ = 20 μM, and C_NMII_ = 0.1 μM.

Video 5. Actin network evolution under d=500nm tensile displacement size, but the time interval between two displacements is reduced from 150s to 10s. The network is first allowed to evolve for 160s before the first pulling event. The video shows the trajectory between 130 ~ 198s with four pulling events in total. The grey sphere represents the pseudo AFM-probe, and red, blue, and green cylinders represent the actin filaments, crosslinkers, and myosin, respectively. C_actin_ = 20 μM, C_NMII_ = 0.1 μM, and C_crosslinkers_ = 2 μM.

**Table 1.**
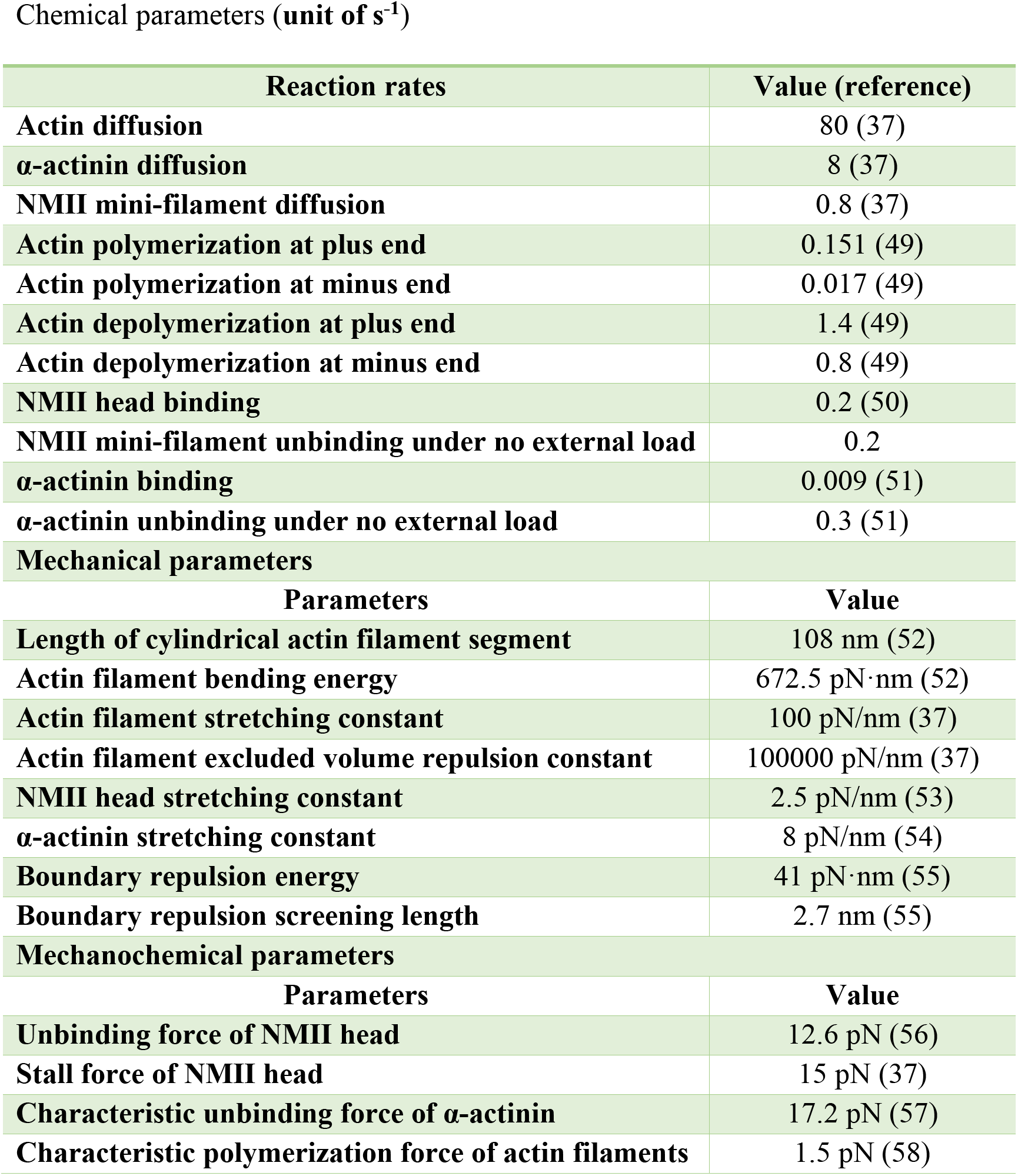
Parameters for the simulations

Chemical parameters (unit of $\text{s}^{-1}$ )
| Reaction rates | Value (reference) |
| --- | --- |
| Actin diffusion | 80 (37) |
| α-actinin diffusion | 8 (37) |
| NMII mini-filament diffusion | 0.8 (37) |
| Actin polymerization at plus end | 0.151 (49) |
| Actin polymerization at minus end | 0.017 (49) |
| Actin depolymerization at plus end | 1.4 (49) |
| Actin depolymerization at minus end | 0.8 (49) |
| NMII head binding | 0.2 (50) |
| NMII mini-filament unbinding under no external load | 0.2 |
| α-actinin binding | 0.009 (51) |
| α-actinin unbinding under no external load | 0.3 (51) |
| **Mechanical parameters** |  |
| Parameters | Value |
| Length of cylindrical actin filament segment | 108 nm (52) |
| Actin filament bending energy | 672.5 pN·nm (52) |
| Actin filament stretching constant | 100 pN/nm (37) |
| Actin filament excluded volume repulsion constant | 100000 pN/nm (37) |
| NMII head stretching constant | 2.5 pN/nm (53) |
| α-actinin stretching constant | 8 pN/nm (54) |
| Boundary repulsion energy | 41 pN·nm (55) |
| Boundary repulsion screening length | 2.7 nm (55) |
| **Mechanochemical parameters** |  |
| Parameters | Value |
| Unbinding force of NMII head | 12.6 pN (56) |
| Stall force of NMII head | 15 pN (37) |
| Characteristic unbinding force of α-actinin | 17.2 pN (57) |
| Characteristic polymerization force of actin filaments | 1.5 pN (58) |

